# Colonization with *Oxalobacter formigenes* slows the progression of CKD and reduces cardiac remodeling in CKD

**DOI:** 10.1101/2025.05.14.654014

**Authors:** Xiaozhong Xiong, Melody Ho, Karim Jaber, Rashmi Mishra, Amalya Charytan, Nadim Zaidan, Florencia Schlamp, Glenn I. Fishman, Lama Nazzal

**Affiliations:** Division of Nephrology, Department of Medicine, NYU Grossman School of Medicine, New York, NY, USA; Leon H. Charney Division of Cardiology, Department of Medicine, NYU Grossman School of Medicine, New York, NY, USA

**Keywords:** Chronic kidney disease, Cardiovascular disease, Hydroxyproline, Microbiome, Oxalate, *Oxalobacter formigenes*

## Abstract

Accumulation of oxalate in patients with chronic kidney disease (CKD) is associated with CKD progression and increased risk of cardiac death. Whether reducing plasma or urine oxalate slows CKD progression and prevents cardiovascular complications remains unexplored. We colonized the intestines of control and CKD mice with *Oxalobacter formigenes* (*Oxf*), an oxalate-degrading microorganism. The mice were fed with the oxalate precursor hydroxyproline for 23 weeks at which time we assessed pathological changes in the kidney and heart. We demonstrate that *Oxf* reduces plasma oxalate (pOx) and creatinine levels, mitigates inflammation and fibrosis in the kidney, and reduces pathologic cardiac remodeling in the hearts of CKD mice. RNA-seq analysis of ventricular tissue of CKD mice reveals dysregulated expression of metabolic pathways while *Oxf* colonization reverses these changes. These findings demonstrate that oxalate accumulation plays a role not only in CKD progression but also in associated cardiovascular complications and suggest that strategies to reduce plasma oxalate levels may have therapeutic benefit.

**Translational statement:** Chronic kidney disease (CKD) is a major health problem that can lead to kidney failure and which increases the risk of cardiovascular disease (CVD) mortality. Oxalate accumulation in advanced kidney disease contributes to further CKD progression and CVD complications. Intestinal colonization with *Oxalobacter formigenes* (*Oxf*) in a CKD animal model reduces plasma oxalate level and slows progression of both CKD and CVD. Strategies to reduce plasma oxalate levels may have therapeutic benefit in the setting of CKD.

## Introduction

Chronic kidney disease (CKD) is a major public health issue affecting 8-16% of the population globally ^1, 2^, often progressing to end-stage kidney disease (ESKD). Patients with CKD face an increased risk of premature cardiovascular (CV) related mortality ^3, 4^. While traditional CV risk factors including hypertension, diabetes and hyperlipidemia contribute to this increased risk, the role of uremic toxins in CVD development and progression has gained increased attention ^2, 5^. Uremic toxins including phosphate, fibroblast growth factor 23 (FGF23), trimethylamine-N-oxide (TMAO), indoxyl sulfate (IS) and p-cresyl sulfate (PCS), have all been implicated in CV complications in CKD ^6–8^. Despite this emerging evidence, mechanisms linking CKD and increased CVD susceptibility remain incompletely understood.

Oxalate, a uremic toxin, is a ubiquitous component of our diet and an end-product of amino acid metabolism which is primarily excreted by the kidneys. In individuals with preserved kidney function, oxalate is efficiently filtered and secreted into the urine. However, as the glomerular filtration rate declines in CKD, oxalate accumulates in the blood leading to various oxalate-related toxicities ^9, 10^. Several epidemiological studies have highlighted the toxic effects of oxalate. In a cohort study of 3,123 patients with CKD, urinary oxalate was identified as an independent risk factor for CKD progression ^11^. Among patients with end stage kidney disease (ESKD) undergoing hemodialysis, higher concentrations of plasma oxalate (POx) have been associated with a higher risk of sudden cardiac death ^3, 12^. These data suggest that diminishing pOx levels may be an important approach to slow CKD progression and mitigate CVD complications.

The gut microbiome has recently emerged as a promising therapeutic target to reduce the burden of CVD in CKD ^13, 14^. Modifying gut microbiota reduced levels of the uremic toxins trimethylamine N-oxide (TMAO) and indoxyl sulfate (IS) thereby reducing the risks of CVD ^15^. Inhibition of microbiota-derived TMAO and IS attenuates CKD progression, prevents cardiac hypertrophy and ameliorates cardiac fibrosis ^16, 17^. Humans lack oxalate-degrading enzymes, making oxalate degradation in the gut entirely reliant on microbial activity ^18, 19^. The “oxalobiome”, or oxalate degrading bacteria, includes several species such as *Oxalobacter formigenes (Oxf*), *Escherichia coli*, *Bifidobacterium* spp., and *Lactobacillus* spp ^18^. Among these, *Oxf* is the most efficient oxalate-degrading bacterium in humans, utilizing oxalate as its sole energy source ^20^. In rodent models, *Oxf* colonization reduced urinary oxalate and calcium oxalate (CaOx) crystal deposition ^21^. Furthermore, in a small uncontrolled trial of primary hyperoxaluria (PH) patients with ESKD, *Oxf* treatment lowered pOx levels and stabilized cardiac function ^22^.

Despite these promising findings, it remains unclear whether *Oxf*-mediated plasma oxalate reduction in CKD contributes to slowing CKD progression and preventing CVD complications. In this study, we investigated the effects *Oxf* intestinal colonization using a CKD mouse model sensitized by exposure to hydroxyproline, an oxalate precursor.

Our results demonstrate that *Oxf* colonization reduces kidney toxicity and mitigates oxalate-mediated cardiovascular remodeling by reducing inflammation and fibrosis in the kidney and heart. Moreover, RNA-seq analysis of heart tissues revealed that *Oxf* colonization ameliorated dysregulation of metabolic pathways in CKD mice. This work provides the first proof that reducing oxalate by *Oxf* colonization is beneficial for CKD induced CVD.

## Materials and Methods

### Animal models

Male C57BL/6 adult mice (40) were purchased from Jackson Laboratories (Bar Harbor, ME). At 6 weeks of age, half underwent two-stage 5/6 nephrectomy (5/6 Nx) surgery, entailing the removal of two-thirds left kidney followed by a right nephrectomy within one week, to induce CKD. The remaining 20 mice underwent sham operations and served as controls. Preoperative care, surgical preparation, and surgical operations were performed by Jackson Laboratories (Bar Harbor, ME) following their standard protocol. Four weeks post-surgery, all animals were transferred to the NYUGSOM animal facility (**Figure 1A**). Animal care and procedures were carried out in compliance with our Institutional Animal Care and Use Committee (IACUC protocol #s16-00822). Mice were housed in groups of five and were maintained on a 12-hour light-dark cycle. They were fed a diet supplemented with 1.5% hydroxyproline and 0.5% calcium (Envigo Teklad Inc., Madison, WI [TD.170633]) for 4 days before transitioning to a diet supplemented with 1% hydroxyproline and 0.5% calcium for the duration of the experiment, as the higher concentration of hydroxyproline induced weight loss. On day 7, the CKD and sham groups were orally gavaged with either one dose of *Oxf* (1×10^9^ CFU in 200µl culture medium) or vehicle (V, 200µl culture medium), resulting in four groups of 10 mice each: CKD+*Oxf*, sham+*Oxf*, CKD+V and sham+V (**Figure 1A**). Intestinal *Oxf* colonization was confirmed via PCR of fecal samples and visualized using gel electrophoresis three days post-gavage and persistent *Oxf* colonization was confirmed using qPCR at weeks 2, 12, and 20 (**Figure 1B**).

**Figure 1.**
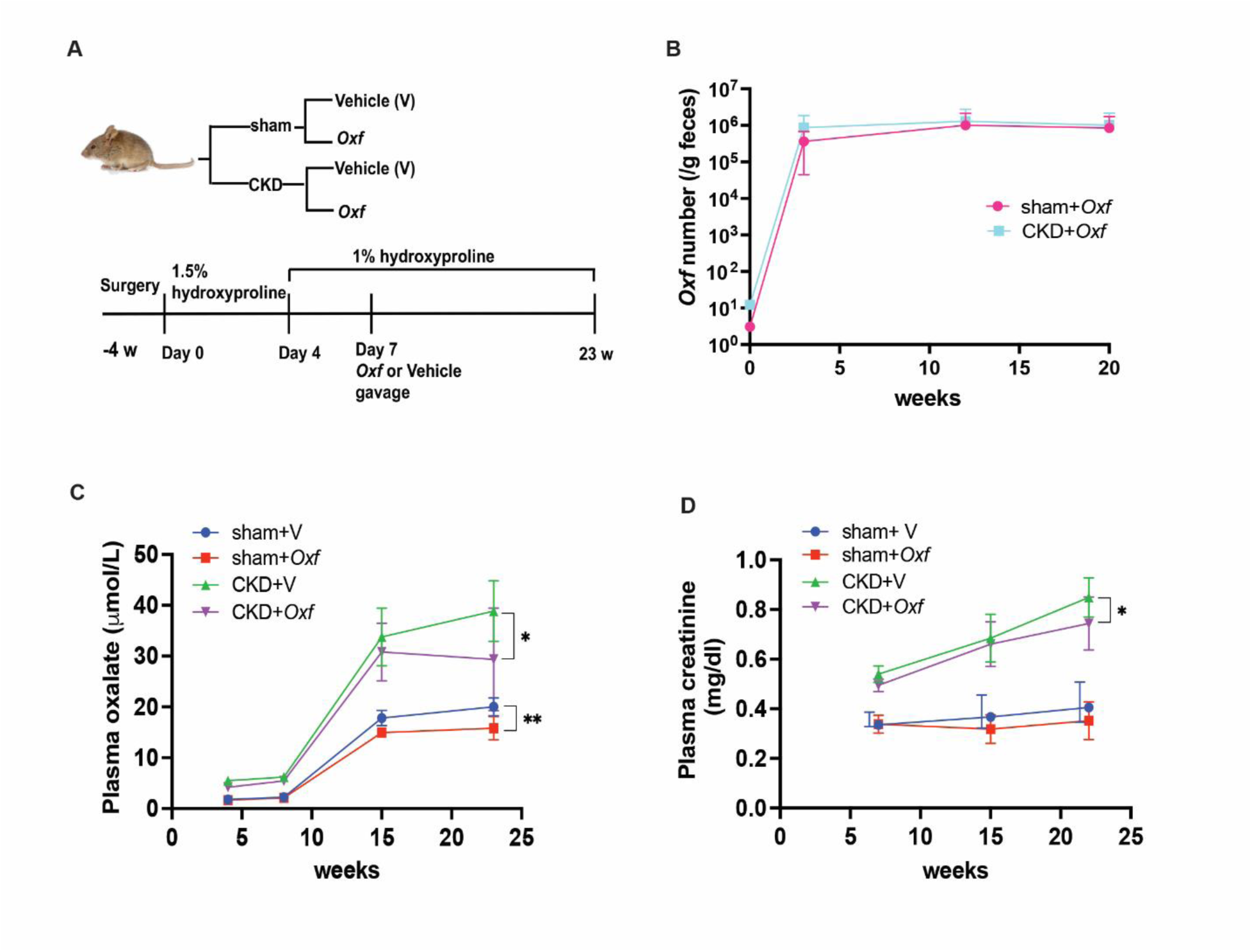
*Oxf* reduces POx and attenuates CKD progression. **A**. Scheme of experiment designation. **B**. *Oxf* was detected by qPCR in mouse feces at different time points after one dose of *Oxf* gavage. **C**. Plasma oxalate level in sham and CKD mice treated with either vehicle (V) or *Oxf*. **D**. Plasma creatinine level in CKD mice treated with either vehicle (V) or *Oxf*. Data present as mean ± SD. * p<0.05; ** p<0.001. T test was performed at week 23 comparing CKD and sham groups to the CKD+*Oxf* and sham+*Oxf* groups, respectively.

The four cohorts were monitored for 23 weeks with weekly fecal collections and biweekly tail-blood collections in EDTA tubes. Plasma was obtained after blood centrifugation for 20 min at 2000 rpm. All samples were frozen and stored at 80°C until use. At the conclusion of the study, all mice were anesthetized with ketamine/xylazine and perfused with PBS at sacrifice. Blood was collected by heart puncture. The heart and kidney were harvested.

### *Oxf* culture

Pure cultures of *Oxf* strain *OxCC13* were grown in a medium consisting of 20mmol/L ammonium oxalate, 40g/L dextrose, 20g/L protease peptone, 10g/L yeast extract, 2ml/L Tween 80, 4g/L KH_2_PO_4_, 10g/L Na-acetate, 4g/L diammonium hydrogen citrate, 0.1g/L MgSO_4_ 7H_2_O and 0.1g/L MnSO_4_. For the determination of colony-forming units (CFU) *OxCC13* was grown in culture media and incubated at 37°C and OD595 measurements were performed using a SpectraMas iD3 plate reader at 24 hours ^23^. Culture at 24 hours was plated on solid plates using the aforementioned medium supplemented with calcium and CFU per milliliter was determined at day 5.

### qPCR for detection of *Oxf* colonization

qPCR was performed to quantitate the copies of *Oxf* strain OxCC13 as previously described ^23^. Fecal DNA was obtained using a Qiagen Powersoil Kit (Hilden, Germany). qPCR was performed using LightCycler 480 SYBR Green І master mix and the LightCycler 480 system (Roche, Basel, Switzerland). The forward primer was 5’-GTGTTGTCGGCATTCCTATC-3 and the reverse primer was 5’-GAAGCAGTTGGTGGTTGC-3’. Cycling conditions included an initial 95°C incubation for 10 minutes, followed by 40 cycles at 95°C for 23 seconds, annealing at 63°C for 20 seconds, and extension at 70°C for 40 seconds.

### Plasma creatinine and oxalate measurement

Plasma creatinine levels were quantified using a Creatinine Serum Detection Kit (Abcam, Cambridge, UK) following the manufacturer’s protocol. Plasma oxalate concentration was measured using the Abcam oxalate kit (Abcam, Cambridge, UK).

### Heart tissue RNA sequencing

Total RNA was extracted from ventricular myocardium using the RNeasy Mini Kit (Qiagen, Hilden, Germany) as per the manufacturer’s protocol while the RNA quality and concentration were assessed using a Nanodrop Spectrophotometer (ThermoFisher Nanodrop 8000 UV-Vis Spectrophotometer). RNA sequencing was performed by the NYU School of Medicine Genome Technology Center on an Illumina Novaseq 6000 instrument. Quality control (QC) was assessed using FastQC (v0.11.7) and MultiQC (v1.9). Reads were mapped to mm10 genome using STAR (v2.6.1d) and genomic features were assigned using Subread featureCounts (v1.6.3). Normalization and differential expression analysis were performed with DESeq2 (v1.30.1). Only genes with average log2 expression > 0.75 in at least 50% of samples were kept. Gene set enrichment analysis (GSEA) was performed on genes with nominal p<0.05 using ClusterProfiler (v3.18.1).

### Histology

The hearts and kidneys were harvested and fixed in 10% formalin. Paraffin-embedded sections were then stained with H&E and Masson trichrome staining (MTS) at the histology core of NYU Langone Medical Center.

Immunohistochemistry (IHC) staining was performed for CD45, alpha-smooth muscle actin (α-SMA), Liver X receptor-alpha (LXRα) and peroxisome proliferator-activated receptor-alpha (PPARα). For antigen retrieval, the paraffin sections were microwave-heated for 10 minutes in 10mM citrate buffer pH 6.0. Endogenous peroxidase activity was blocked with 0.3% hydrogen peroxidase in PBS for 20 min, after which slides were washed and incubated with 5% normal goat serum in PBS for 1 hour at room temperature. Primary antibodies include rabbit anti-CD45 monoclonal antibody, rabbit anti-α-SMA monoclonal antibody (Cell Signaling Technology, Danvers, MA), rabbit anti-LXRα polyclonal antibody (LifeSpan Biosciences, Lynnwood, WA) and rabbit anti-PPARα (Thermo Fisher Scientific, Waltham, MA). Sections were incubated with primary antibody overnight at 4°C followed by incubation with secondary HRP-conjugated goat anti-rabbit IgG (Thermo Fisher Scientific, Waltham, MA) for 1h. For LXRα and PPARα staining, biotin-conjugated goat anti-rabbit IgG and ABC kit (Vector Laboratories, Newark, CA) were used. A DAB (3,3’-diaminobenzidine) substrate kit (Sigma, Louis and Burlington, MA) was used for detecting HRP activity.

### Laser scanning confocal microscopy

Paraffin embedded heart tissues were stained with Wheat Germ Agglutinin (WGA) Alexa Fluor™ 488 Conjugate (50μg/ml, W11261, Thermo Fisher Scientific, Waltham, MA) at room temperature for 1h and washed with PBS four times. Tissue sections were imaged using a confocal fluorescence microscope (Zeiss, LSM 800).

### Transmission Electron Microscopy

A piece of the ventricle was cut from paraffin-embedded mouse heart. The heart was deparaffinized using xylene, and rehydrated to ddH_2_O via a gradient of ethanol (100%, 95%, 85%, 70%, 50%, 30%, 10 minutes each step). After washing the tissue with PBS (pH 7.4) for 25 minutes, the tissue was fixed in 2.5% glutaraldehyde in 0.1 M phosphate buffer (pH 7.4) for 2 hours. The samples were washed 3 times with 0.1 M phosphate buffer (10 minutes each time), post-fixed with 1% osmium tetroxide with 1% potassium ferrocyanide for 1.5 hours on ice and dehydrated in a serial of ethanol solutions (30, 50, 70, 85, 95, 100%; 10 min each). The heart tissues were then washed in propylene oxide twice for 5 minutes each, infiltrated with propylene oxide and embedded in EMbed 812 (Electron Microscopy Sciences, Hatfield, PA). Semi-thin sections were cut at 500 nm and stained with 1% toluidine blue to find the orientation of the heart tissue. Ultrathin sections of 70 nm were cut, mounted on 200 mesh copper grids, stained with uranyl acetate and lead citrate by standard methods, and imaged with JEOL 1400 Flash transmission electron microscope (Japan) using Gatan Rio16 camera (Gatan, Inc., Pleasanton, CA).

### Quantification and Statistics

The percentage of CD45+ cells, indicating the number of inflammatory cells, was quantitated by counting CD45+ cells and total cells in 8-10 random fields from each slide and dividing CD45+ cells by total cells. α-SMA positive area and trichrome Masson staining (TMS) positive area were measured by Image J. Perivascular collagen and blood vessel wall were measured by ImageJ and perivascular fibrosis area determined by ratio of collagen in adventitial area and thickness of blood vessel wall ^24^. Slides were from 4-6 mice of each group and 8-10 random fields of each slide were measured.

Cardiomyocyte area was quantified from WGA-stained heart sections using ImageJ, analyzing 4–5 cross-sectional fields from 3–4 mice per group.

Data were analyzed using GraphPad Prism 9.2. Values are presented as mean ± standard deviation. Statistical differences between grouped quantitative datasets were measured using the 2-tailed unpaired Student’s test. A p-value of < 0.05 is considered statistically significant.

For longitudinal plasma creatinine and oxalate analysis, an analysis of variance (ANOVA) was conducted to assess the differences in plasma (oxalate or creatinine) area across treatment groups.

The treatment group was used as the independent variable, and the plasma area was treated as the dependent variable. To investigate the specific differences between treatment groups, a Tukey Honest Significant Difference (HSD) test was performed following the ANOVA. The adjusted p-values and the corresponding confidence intervals were calculated to determine which pairwise differences are statistically significant.

## Results

### *Oxf* intestinal colonization decreases plasma oxalate levels and attenuates CKD progression

We first confirmed, using qPCR, that a single oral gavage of *Oxf* is sufficient for long-term intestinal colonization in our mouse model (**Figure 1B**). pOx levels were first measured after 4 weeks of hydroxyproline diet, and as expected, we observed a higher pOx in CKD+V mice than sham+V mice. pOx levels increased over time in both sham+V and CKD+V mice, with CKD+V mice showing consistently higher levels than sham mice over time. *Oxf* colonization resulted in lower pOx levels over time in CKD+*Oxf* mice but not in sham+*Oxf* mice, compared with CKD+V and sham+V, respectively (**Figure 1C**). At sacrifice, pOx was 21% lower in sham+*Oxf* mice compared to sham+V mice (15.8±2.28 μM/L vs 20.01±1.76 μM/L, p=0.0004), and 24% lower in CKD+*Oxf* compared to CKD+V mice (29.35±10.06 μM/L vs 38.85±5.96 μM/L, p=0.038). To assess kidney function, plasma creatinine (pCr) was first measured 7 weeks following the start of the diet and CKD+V mice exhibited significantly higher pCr levels than sham+V mice (**Figure 1D).** pCr levels gradually increased over time in all CKD+V mice, but at sacrifice, CKD+*Oxf* mice had significantly lower pCr levels than CKD+V mice (0.74±0.10 mg/ml vs 0.85±0.08 mg/ml, p=0.037). Sham+V and sham+*Oxf* mice had comparable pCr at sacrifice. However, over the period of the experiment, pCr was not significantly different in the CKD+*Oxf* and sham+*Oxf* as compared to their respective controls (**Figure 1D)**. These results indicate that Ox*f* colonization significantly decreases plasma oxalate in CKD mice and reduces progression of CKD.

### *Oxf* impedes inflammation and fibrosis in the kidney

CKD is characterized by progressive and irreversible nephron loss, microvascular damage, inflammation, and fibrosis, ultimately leading to renal failure and ESKD ^25^. Oxalate crystals (CaOX) are associated with renal inflammation, fibrosis, and progressive renal failure ^26^. We observed that lowering pOx was associated with a slower rate of creatinine rise in CKD mice. To determine whether reduction in pOx by *Oxf* gut colonization mitigates the pathological progression of CKD, we examined inflammation and fibrosis in mouse kidneys using IHC staining for the inflammatory cell marker CD45, fibroblast marker α-SMA, and Masson’s trichrome staining (MTS). As shown in Figure 2, CD45+ cells were present in the kidneys of both CKD+V and sham+V mice, but the infiltration of CD45+ cells was greater in CKD+V mice (**Figure 2A and 2E**). Moreover, *Oxf*-colonization led to a lower CD45+ cell infiltration in the kidney of both sham+*Oxf* and CKD+*Oxf* groups (**Figure 2A and 2E**). *Oxf*-colonized sham and CKD mice have a substantially smaller kidney fibrosis area than their control sham+V and CKD+V mice (**Figure 2B and 2F**). Further, evidence from α-SMA staining suggests a markedly lower fibrotic area in the kidneys of sham+*Oxf* and CKD+*Oxf* mice than sham+V and CKD+V mice (**Figure 2C and 2G**), respectively, consistent with the MTS staining results. These results indicate that *Oxf* gut colonization attenuated kidney inflammation and fibrosis in both sham and CKD mice, and the reduction of inflammation and fibrosis was greater in the CKD groups compared to the sham groups. To further assess kidney pathology and CaOx crystal deposition, we performed Periodic Acid-Schiff (PAS) staining and Pizzolato staining, respectively. Compared to sham+V mice, CKD+V mice kidneys exhibited enlarged tubules and irregular as well as thicker glomerular membranes, indicating severe kidney injury in CKD mice (**Figure 2D**). While *Oxf*-colonized CKD mice also exhibited enlarged tubules and irregular and thicker glomerular membranes, they had less severe glomerular and tubular damage compared to CKD+V mice (**Figure 2D**). Pizzolato staining showed that CaOx crystals were deposited in CKD+V mice but not in sham+V mice; *Oxf* treated CKD mice had fewer CaOx crystals than CKD+ V mice (**Figure S1**).

**Figure 2.**
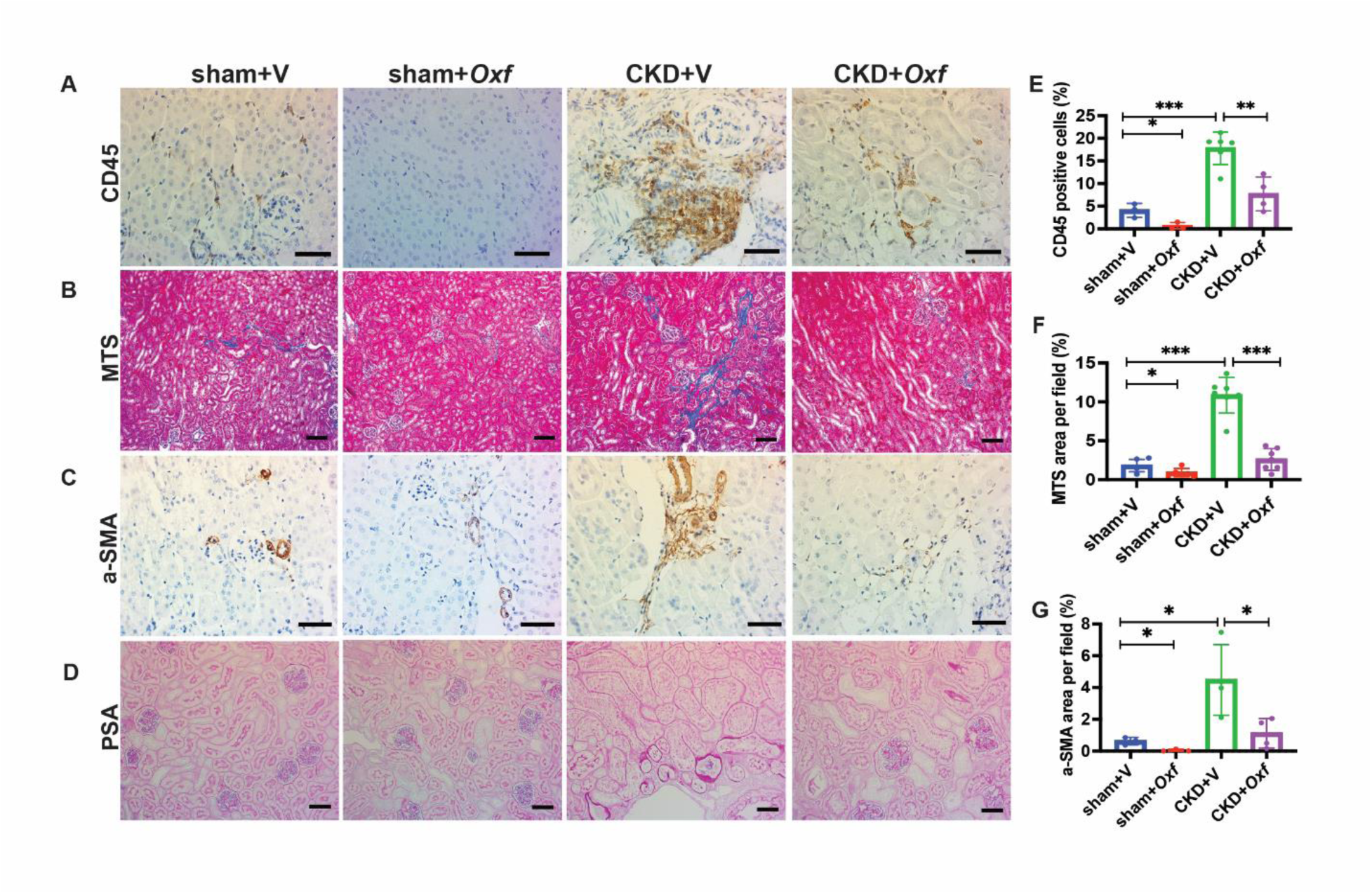
Histological assessment of inflammation and fibrosis in the kidneys. **A**. Representative images of CD45 staining. Scale bar = 50 mm. **B**. Representative images of Manson Trichrome Staining (MTS). Scale bar = 100 mm. **C**. Representative images of a-SMA staining. Scale bar = 50 mm. **D**. Representative images of Periodic Acid Schiff (PSA). Scale bar = 50 mm. **E.** Kidney tissues were stained with CD45 and percentage of CD45+ cells were calculated in sham and CKD mice treated with either vehicle (V) or *Oxf*. **F**. Manson’s Trichrome Staining (MTS) in kidney tissues and percentage of MTS positive area per field were measured in sham and CKD mice treated with either vehicle (V) or *Oxf.* **G**. a-SMA staining in kidney tissues and percentage of a-SMA positive area per field were calculated in sham and CKD mice treated with either vehicle (V) or *Oxf*. Data presented as mean ± SD. * p<0.05; ** p<0.01; *** p<0.001.

### *Oxf* reduces fibrosis and angiogenesis in the heart of CKD

Patients with CKD exhibit characteristic changes in the myocardium with pathological myocardial fibrosis with collagen deposition and cardiac hypertrophy, the hallmarks of CKD induced cardiomyopathy ^3, 27^. Since the impact of oxalate on cardiac histological remodeling has not been previously evaluated, we performed α-SMA and MTS staining on ventricular tissue to assess angiogenesis and fibrosis, respectively, and performed WGA staining to evaluate cardiac hypertrophy. As expected, the number of α-SMA expressing blood vessels in the ventricles of CKD+V mice was significantly higher compared to sham+V mice (24.88 ± 4.92/mm^2^ vs. 14.06 ± 1.71/mm^2^, p=0.0004, **Figure 3A and 3F**), indicating increased angiogenesis. However, CKD+*Oxf* mice had significantly lower number of blood vessels than CKD+V mice (17.05±2.95/mm^2^ vs. 24.88±4.92/mm^2^, p=0.023, **Figure 3A and 3F**). There was no significant difference in the number of blood vessels between sham+V and sham+*Oxf* mice (**Figure 3A and 3F**). In CKD+V mice, α-SMA was expressed in cardiomyocytes and pericytes (**Figure 3A and 3B, arrows indicated**) but was confined to the smooth muscle layers in all sham (+V or +Oxf) and CKD+*Oxf* mice. Perivascular fibrosis was illustrated by an increase in perivascular collagen deposition stained with MTS. Increased perivascular collagen deposition was observed in the hearts of CKD+V mice (**Figure 3C, arrow indicated**) but not in sham+V mice (**Figure 3C**) and CKD+*Oxf* mice had less collagen deposition in their perivascular area than CKD+V mice (p=0.006) (**Figure 3G**).

**Figure 3.**
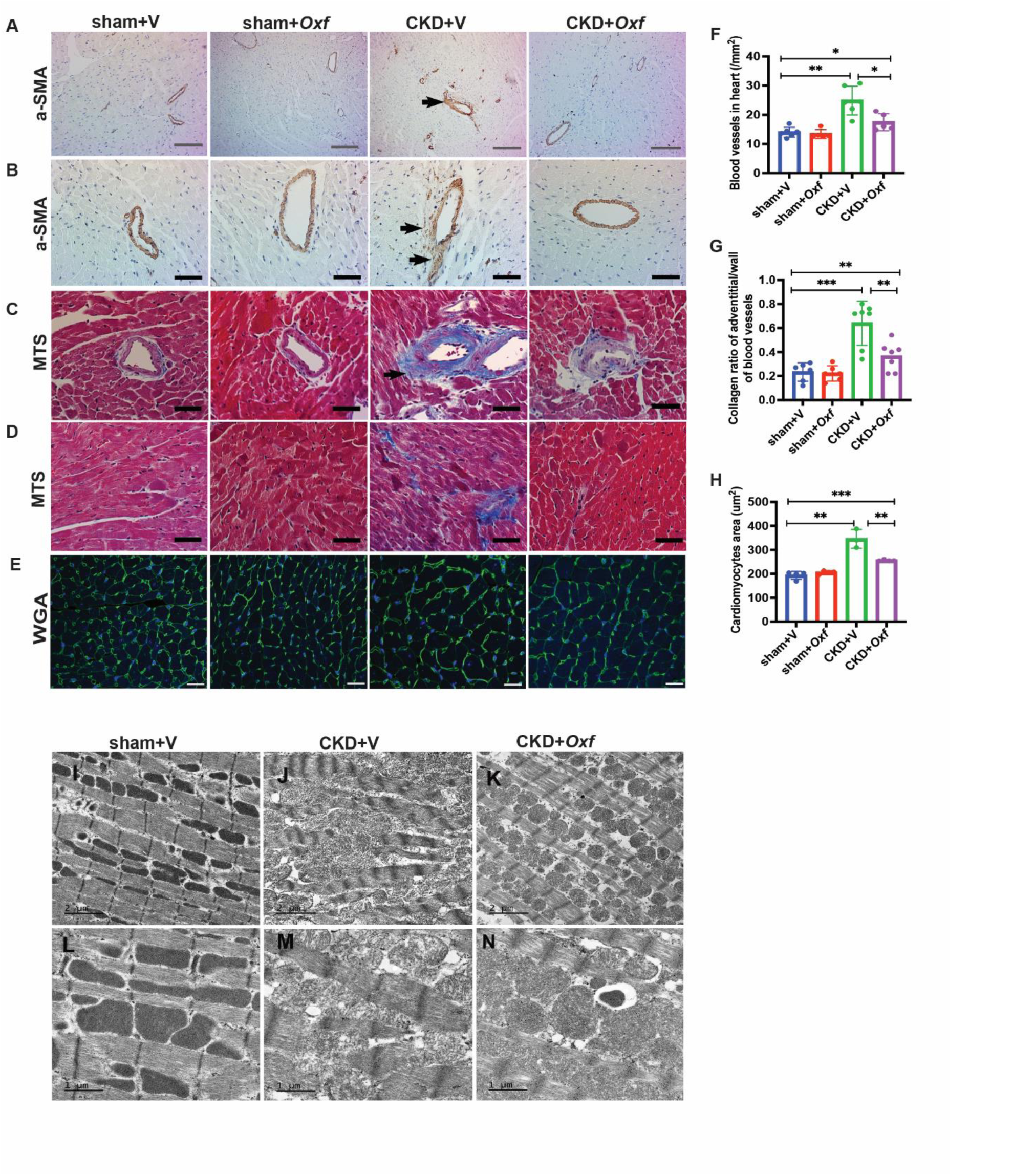
Histological assessment of angiogenesis, fibrosis and mitochondrial imaging in the heart. **A**. Representative images of over all look of blood vessels in different groups of mice. Scale bar = 200 mm. **B.** Representative images of blood vessels in higher magnification. Arrows indicate perivascular fibrosis. Scale bar = 50 mm. **C**. Representative images of collagen deposition in perivascular area in different groups of mice. Scale bar = 50 mm. **D**. Representative images of collagen deposition in interstitial area in different groups of mice. Scale bar = 50 mm. **E**. Representative images of WGA staining in the heart tissues. Scale bar = 20 mm. **F**. Heart tissues were stained for a-SMA. a-SMA positive blood vessels in ventricles were measured and calculated by Image J. Data present as mean ± SD. **G**. Heart tissues were stained for MTS. Collagen deposition in perivascular area was measured and calculated by Image J. Data present as mean ± SD. **H**. Heart tissues were stained for WGA. The area of cardiomyocytes was measured by Image J. Data present as mean ± SD. **I-N**. Mitochondria morphology of ventricle tissues was examined by TEM. Representative images of mitochondria in sham+V, CKD+V and CKD+*Oxf* mice. **I-K:** Scale bar = 2 mm. **L-N:** scale bar = 1mm. * p<0.05; ** p<0.01; *** p<0.001

Interstitial fibrosis was only found in 4 out of 8 CKD+V mice (**Figure 3D**) while absent in all other groups. WGA staining demonstrated that the size of cardiomyocytes in the heart of both CKD+V and CKD+*Oxf* mice were significantly larger compared to sham+V mice (**Figure 3E and 3H**). However, cardiomyocytes were significantly smaller in CKD+*Oxf* mice compared to CKD+V mice (**Figure 3E and 3H**). These results indicate that CKD mice had evidence of coronary angiogenesis, perivascular fibrosis, and cardiac hypertrophy, while *Oxf* colonization reduced the extent of pathologic cardiac remodeling in CKD mice.

### *Oxf* reduces mitochondrial morphological abnormalities in the heart of mice with CKD

To further investigate CVD complications, we assessed the effect of high oxalate diet on cardiac mitochondria. Mitochondrial dysfunction plays a key role in cardiovascular disease, particularly cardiac hypertrophy and heart failure ^28, 29^, due to the high mitochondria content and oxygen consumption of the heart. We examined mitochondrial morphology of ventricular tissues of sham+V, CKD+V and CKD+*Oxf* mice using transmission electron microscopy (TEM). Mitochondria in the heart of sham+V mice showed normal features: mitochondria were aligned and well organized with intact membranes and tightly packed cristae. Z lines were linear, myofibrils were parallel and well aligned **(Figure 3I** and **3L).** In the CKD+V mice, mitochondria were swollen and disorganized with loose and enlarged cristae, Z lines were waved, and myofibrils were disrupted **(Figure 3J and 3M).** However, compared to CKD+V mice, mitochondria in CKD+Ox*f* mice exhibited less swelling and had better alignment and organization. Mitochondria from CKD+*Oxf* mice had tightly packed cristae, neatly preserved membranes and well-arranged myofibrils (**Figure 3K and 3N**). These results demonstrate that CKD+V mice having more severe mitochondrial damage than *Oxf* treated CKD mice.

### RNA-seq analysis reveals dysregulation of metabolic pathways in the heart of CKD mice

To gain mechanistic insight into the pathogenesis of heart disease in CKD and explore how reduction of oxalate using *Oxf* ameliorates disease severity, we performed RNA-seq on ventricular tissue from each of the four cohorts of mice. We first analyzed differential gene expression in CKD+V vs sham+V mice and identified 87 differentially expressed transcripts (P_adj_ <0.05). Out of these 87 genes, 24 were up-regulated and 63 were down-regulated in CKD+V mice (**Figure 4A and 4B**). Top upregulated genes (fold>2.0, p _adj_ <0.01) in CKD+V were Deleted in malignant brain tumors 1 (Dmbt1) and Carboxypeptidases A1 (Cpa1) (**Figure 4C**). Top downregulated genes (fold>2.0, p _adj_ <0.01) in CKD+V included members of the Major urinary protein (Mup) family, especially Mup14, Mup8 and Mup1 (**Figure 4D**). Genes related to oxidative phosphorylation (OXPHOS) (**Figure 4E**) and cardiomyopathy were upregulated in CKD+V mice (**Figure 4F**). Upregulation of OXPHOS genes suggests an increase in energy demand in the hearts of CKD+V mice. Interestingly, fetal genes Myh7 and Nppa were also increased in the hearts of CKD mice (**Figure 4F**). Increase of fetal gene expression in the adult heart is a hallmark of several forms of pathologic remodeling ^30^. Gene Ontology (GO) enrichment analysis revealed that the changes in gene sets were positively associated with cardiovascular disease and mitochondrial dysfunction in CKD+V, some of which included left ventricular hypertrophy, mitochondrial respiratory chain complex, and regulation of cardiac muscle contraction (**Figure 4G**). Ingenuity Pathways Analysis (IPA) revealed that LXR/RXR activation, DHCR24 signaling pathway and regulation of lipid metabolism by PPARα were significantly inhibited, while IL-12 signaling was significantly upregulated in CKD+V (**Figure 4H**). LXRs, DHCR24, and PPARs are known to regulate cholesterol and lipid metabolism, mitochondrial function and inflammation ^31–34^. These results suggest that altered genes and pathways in CKD are related to cardiomyopathy as well as mitochondrial dysfunction.

**Figure 4.**
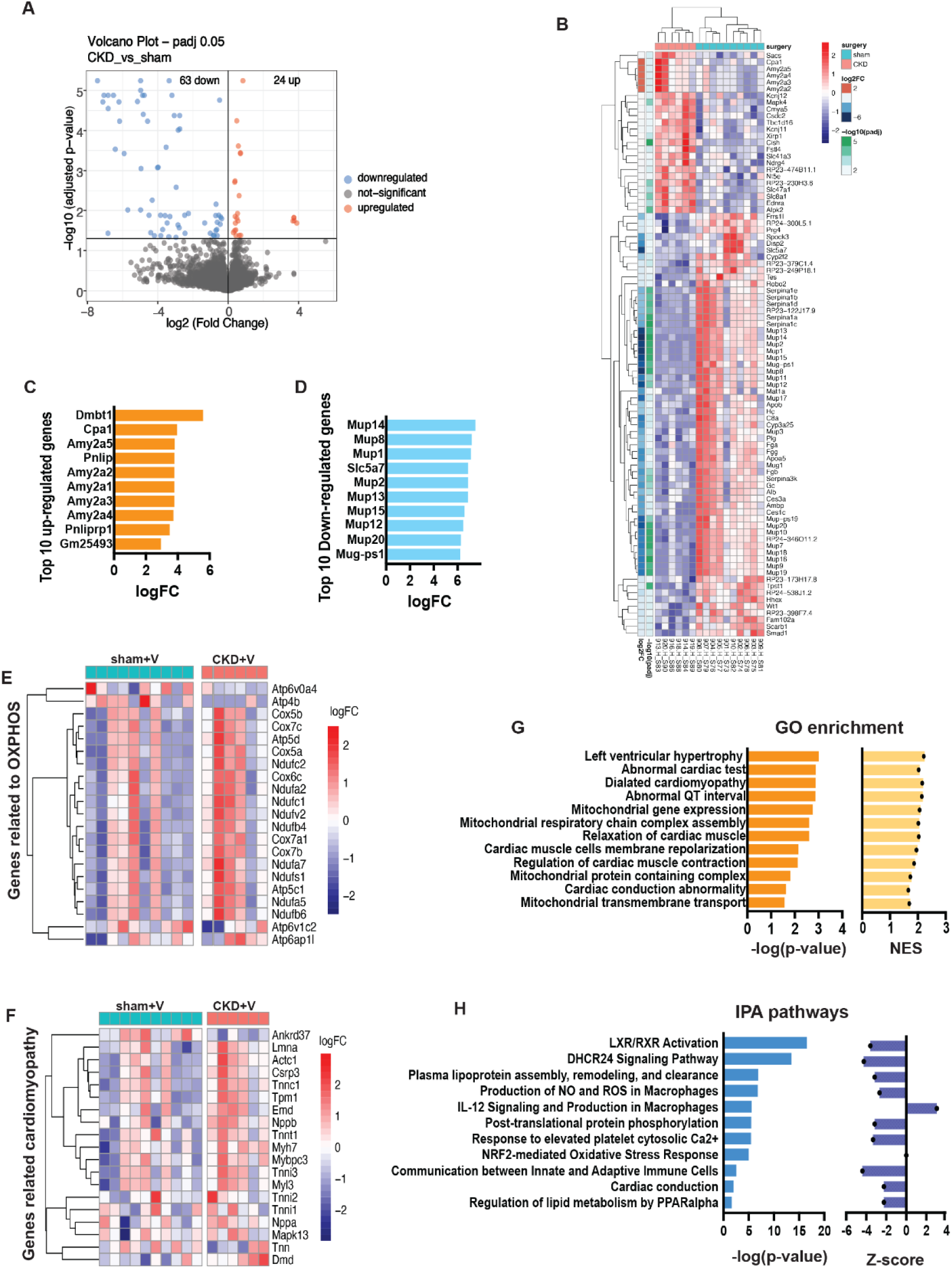
RNA-seq analysis of genes and pathways in the hearts of in CKD compared to sham mice. **A**. Volcano plot displayed differentially expressed genes (adjusted p<0.05). **B**. Heatmap of differentially expressed genes (adjusted p<0.05). **C**. Top 10 up-regulated genes (fold >2, p<0.01). **D**. Top 10 down-regulated genes (fold >2, p<0.01). **E**. Heatmap of differentially expressed genes related to oxidative phosphorylation (OXPHOS) (p<0.05). The color intensity represents log2FC changes. **F**. Heatmap of differentially expressed genes related to cardiomyopathy (p<0.05). The color intensity represents log2FC changes. **G**. GO enrichment of the significantly changed cellular components of the heart. Right side (orange) is -log(p-value), left side (yellow) is NES value. **H**. IPA pathways of differentially expressed genes (DEG) of the heart. Right side (blue) is -log(p-value), left side (dark purple) is Z-score score.

### *Oxf* treatment reverses genes and pathways altered in CKD

We next analyzed differential gene expression in CKD+*Oxf* mice vs CKD+V mice. There were 236 genes differentially expressed in CKD+*Oxf* mice compared to CKD+V mice (125 genes upregulated, and 111 genes downregulated (P_adj_ <0.05) (**Figure S2A and S2B**). Interestingly, we found that the top altered genes in CKD+V were reversed by *Oxf* colonization in CKD+*Oxf* (**Figure 5A).** The most highly upregulated genes in CKD+V group were significantly downregulated in CKD+*Oxf* group, and vice versa (**Figure 5B, Figure S2C and S2D)**. Genes related to cardiomyopathy and mitochondrial dysfunction were also reversed in CKD+*Oxf* (**Figure 5C and 5D, Figure S2E and S2F**). Notably, fetal genes Nppa and Nppb were downregulated in CKD+*Oxf* compared to CKD+V control, suggesting that *Oxf* reversed fetal gene programming in the heart of mice with CKD thereby reducing the severity of CVD. Similar to the gene pattern, the most altered pathways by GO enrichment analysis and IPA analysis in CKD+V were also reversed in CKD+*Oxf*. GO enrichment analysis showed that the pathways in CKD+*Oxf* mice had a negative association with cardiomyopathy (**Figure 5E, Figure S2G and S2H**). IPA analysis showed that LXR/RXR activation pathway, PPARα pathway and DHCR24 signaling were upregulated, while IL-12 signaling was downregulated in CKD+*Oxf* mice (**Figure 5F**). These results indicate that *Oxf* colonization reversed the CVD-related transcriptional changes in CKD mice and suggest that *Oxf* colonization prevented the dysregulation of CVD-related signaling.

**Figure 5.**
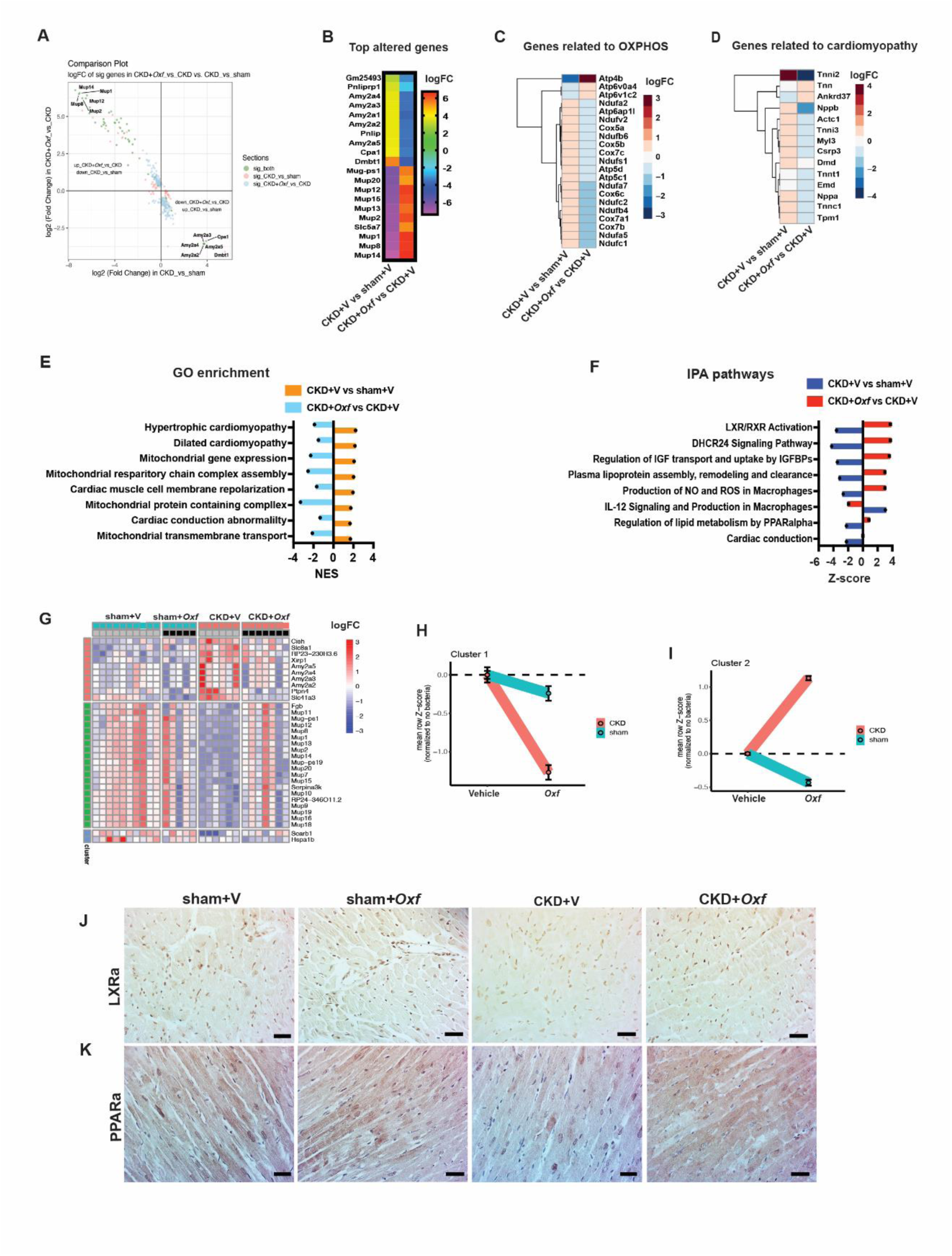
RNA-seq analysis of genes and pathways in the hearts of CKD+*Oxf* vs CKD+V compared to CKD+V vs sham+V. **A**. Volcano plot displayed differently expressed genes. **B**. Top 10 upregulated and top 10 downregulated genes. **C**. Comparison of CKD+V vs sham+V and CKD+*Oxf* vs CKD+V on the gene expression of oxidative phosphorylation (OXPHOS) (P<0.05). The color intensity represents log2FC changes. **D**. Comparison of CKD+V vs sham+V and CKD+*Oxf* vs CKD+V on the gene expression of cardiomyopathy (p<0.05). The color intensity represents log2FC changes. **E**. Comparison of CKD+V vs sham+V and CKD+*Oxf* vs CKD+V on the significantly changed cellular components in GO enrichment. Light blue color indicates down-regulation, orange color indicates up-regulation. **F**. Comparison of CKD+V vs sham+V and CKD+*Oxf* vs CKD+V on the IPA pathways. Blue color indicates inhibition, red color indicates activation. **G**. Linear regression analysis revealed 32 genes were significantly expressed (adjusted p<0.05) due to the effects of surgery (CKD) alone, and hierarchical clustering on the expression of these genes showed 3 gene clusters. **H**. Effect of *Oxf* treatment on gene cluster 1 in sham and CKD. **I**. Effect of *Oxf* treatment on gene cluster 2 in sham and CKD. **J**. Representative images of LXRa staining in the heart tissues of different groups of mice. Scale bar = 50 mm. **K**. Representative images of PPARa staining in the heart tissues of different groups of mice. Scale bar = 50 mm.

To further understand the extent to which both the CKD phenotype and *Oxf* colonization affected the transcriptional responses, we performed linear regression analysis to model the effects of surgery (Sham/CKD), bacterial colonization (*Oxf/*vehicle), and their interactions. This analysis identified 32 genes significantly altered by surgery alone (p_adj_ <0.05), and hierarchical clustering of these genes revealed three distinct clusters **(Figure 5G).** Genes in cluster 1 were upregulated while genes in cluster 2 were downregulated relative to sham+V and sham+*Oxf* groups (**Figure 5G**), suggesting these alterations were caused by CKD. Cluster 1 genes are associated with inflammation and cardiovascular diseases ^35–37^, whereas cluster 2 genes are linked to lipid metabolism and mitochondrial function ^38, 39^. Notably, *Oxf* colonization reversed these alterations in both clusters in CKD mice **(Figure 5G).** In contrast, *Oxf* colonization had no significant effect on the expression of clusters 1 and 2 in sham mice (**Figure 5H and 5I),** indicating its modulatory role is specific to the CKD context.

Given that the LXR activation pathway and PPARα signaling pathway were significantly altered in our RNA-seq analysis, and considering the critical roles of LXRα and PPARα in CVD ^33, 40^, we further evaluated their expression in the heart using IHC staining. Both LXRα and PPARα expression levels were reduced in the hearts of CKD+V mice compared to sham+V mice. However, *CKD+Oxf* mice exhibited expression of LXRα and PPARα at levels comparable to those inham+V mice (**Figure 5J and 5H**). There was no difference in LXRα and PPARα expression between sham+V and sham+*Oxf* (**Figure 5J and 5H**). These findings indicate that LXRα and PPARα are downregulated in the CKD heart, and that *Oxf* colonization can restore their expression.

## Discussion

CVD remains the leading cause of death in CKD patients and represents a persistent clinical challenge. While traditional CVD treatments are effective in the general population, they are only partially successful in reducing mortality rates among CKD patients ^3, 41^. Therefore, identifying non-traditional risk factors that contribute to this elevated risk is a critical research priority. Among these, elevated pOx levels— commonly observed in CKD patients—have drawn increasing attention for their potential role in CVD complications.

Oxalate has been implicated in adverse CV outcomes by inducing oxidative stress in blood vessels and exacerbating atherosclerosis ^42^. In mice, a high-oxalate diet has been shown to induce CKD, accompanied by arterial hypertension and cardiac fibrosis ^43, 44^. Cardiac remodeling, a hallmark of CVD, is characterized by coronary angiogenesis, fibrosis, and mitochondrial dysfunction, all of which contribute to cardiac hypertrophy and progression to heart failure ^45–49^. However, the mechanisms underlying oxalate-induced cardiotoxicity remain poorly understood. Despite its clinical significance, no approved therapies currently exist to efficiently reduce pOx levels in CKD patients.

In our study, CKD mice exhibited pathologic cardiac remodeling, including perivascular and interstitial fibrosis, increased cardiac angiogenesis and cardiac hypertrophy, consistent with previous findings ^3, 27, 45^. The myocardium’s secretion of angiogenic growth factors drives vascular growth to meet the increased metabolic demands, promoting myocardial hypertrophy, even in the absence of direct hypertrophic stimuli ^50^. *Oxf* intestinal colonization mitigated CKD-induced cardiac remodeling as evidenced by reduced coronary angiogenesis, perivascular fibrosis, mitochondrial damage and cardiac hypertrophy. These findings suggest that these pathological changes are mediated, at least in part, by the uremic toxin oxalate. The protective effect of *Oxf* on cardiac remodeling in CKD is likely attributable to both direct inhibition of cardiac toxicity as well as indirectly through slowing of CKD progression.

The molecular mechanisms of oxalate toxicity in cardiomyocytes remain poorly understood. However, oxalate-induced mitochondrial dysfunction and oxidative stress have been observed in endothelial cells, monocytes, macrophages, and renal epithelial cells ^42, 51–53^. To explore how *Oxf* protects against cardiac remodeling in CKD, we performed RNA -sequencing on ventricular tissues from mice. This analysis revealed significant alterations in genes and pathways associated with cardiomyopathy, lipid metabolism dysfunction, inflammation, and mitochondrial dysfunction in CKD mice compared to sham mice.

Key findings included the upregulation of Dmbt1 and Cpa1, which are implicated in angiogenesis and vascular diseases, and the downregulation of the Mup family, associated with lipid metabolism dysfunction and mitochondrial dysfunction ^38, 54–56^. Genes related to OXPHOS and cardiomyopathy were also upregulated, reflecting increased metabolic demands in the heart possibly as a compensatory response to hypertrophic cardiomyocytes and heart failure ^57–59^. The upregulation of fetal genes Myh7 and Nppa provided further evidence of a cardiomyopathic state in the CKD mice.

Notably, *Oxf* treatment reversed the dysregulation of these genes and pathways, highlighting its ability to improve mitochondrial function and prevent cardiac remodeling. While there are numerous uremic toxins that accumulate in the setting of CKD, our findings suggest that oxalate itself is sufficient to promote cardiac disease progression. Moreover, our data indicate that reducing pOx by *Oxf* colonization may serve as a novel and relatively simple therapeutic strategy to blunt pathologic cardiac remodeling.

Interestingly, the LXR activation pathway and PPARα signaling pathway were significantly downregulated in CKD mice and these results were corroborated by immunostaining studies showing decreased LXRα and PPARa protein abundance in CKD hearts. LXRs and PPARs are key regulators of cardiac metabolism and play central roles in cardiac disease ^33, 60^. LXRs regulate cholesterol metabolism, fatty acid catabolism, β-oxidation, inflammation, and oxidative stress ^43, 61–63^. Inhibition of LXRs leads to lipid accumulation, mitochondrial dysfunction, and inflammation, all of which contribute to the progressive cardiac dysfunction. Similarly, PPARs regulate fatty acid oxidation, fat cell development, lipoprotein metabolism, and glucose homeostasis ^60, 61^. PPARs activation increases fatty acid uptake and enhances ATP production, whereas its inhibition disrupts mitochondrial function and leads to metabolic disorders ^33^.

Our results are consistent with studies linking oxalate to alterations in the LXRs and PPARs in other organs. Downregulation of LXRs has been observed in kidney tissues from human CaOx stone formers, and in a CaOx stone murine model ^62^. Renal tubular epithelial cells treated with CaOx crystals show downregulation of LXRs, resulting in increased production of ROS and apoptosis ^62^. Additionally, oxalate treatment in hepatocytes has been shown to suppress PPARα activity, resulting in lipid accumulation and reduced fatty acid oxidation ^63^.

In our study, the inhibition of these pathways in CKD mice is likely driven by elevated pOx levels. Conversely, their upregulation in *Oxf*-colonized CKD mice suggests that reducing oxalate toxicity through *Oxf* colonization can restore these critical metabolic pathways. Further studies are needed to confirm the mechanism linking pOx to cellular metabolic alterations and mitochondrial dysfunction and to assess their impact on cardiac contractile and electrophysiological physiology.

While a high oxalate diet is known to cause oxalate nephropathy in humans and induce CKD in rodent models ^25, 43, 64^, primarily through CaOx crystal deposition and subsequent renal inflammation ^26^, our study is the first to evaluate the impact of gut colonization with *Oxf* on CKD progression. This work addresses a critical gap in understanding how microbiome modulation can influence systemic oxalate levels and their downstream effects on the kidneys. We demonstrated that modifying the gut microbiome through *Oxf* colonization significantly lowers pOx levels and has potential to slow CKD progression.

Histological assessments revealed that *Oxf* colonization prevented renal fibrosis and inflammation in both CKD and sham mice, reinforcing the therapeutic potential of reducing oxalate as a treatment strategy for oxalate nephropathy. Multiple studies have shown that calcium oxalate crystals induce NLRP3 inflammasomes activation, contributing to inflammation ^65^. Our findings are further supported by the lower prevalence of CaOx crystals in *Oxf* treated mice compared to CKD controls.

Interestingly, sham mice, which did not have CaOx crystals, also demonstrated protective effects from *Oxf* colonization. This suggests that *Oxf’s* benefits extend beyond crystal reduction, potentially through crystal-independent mechanisms that mitigate oxalate nephrotoxicity.

The use of oxalate-supplemented diet has been a common method to induce hyperoxalemia and CKD in murine models ^44^. However, our study demonstrates that hyperoxalemia can also be induced by supplementation with hydroxyproline, an oxalate precursor. Hydroxyproline is present in collagen, making meat a significant source of this compound. This connection suggests that hydroxyproline may contribute to the association between high animal protein diets and poor renal and cardiovascular outcomes in CKD ^66^. Additionally, our data highlights the need for caution regarding the use of collagen supplements in this population, as they could conceivably exacerbate hyperoxalemia and its associated complications.

While our study provides strong preclinical evidence, several limitations should be addressed in future research. Notably, we focus on cardiac structure and gene expression, but did not directly assess cardiac function. Second, the potential impact of *Oxf* on the gut microbiome was not studied. However, given the low abundance of *Oxf* in the gut, significant effects on the microbiome’s community structure are unlikely.

In summary, our findings demonstrate that colonization with oxalate-degrading bacteria in the setting of CKD alleviates pathologic cardiac remodeling. Future epidemiological and interventional studies are crucial for translating these findings into clinical practice and exploring broader applications of gut microbiome modulation in CKD and CVD management.

## Supplementary Material

Supplementary Figures S1 and S2 (pdf)

## Acknowledgment

This work is supported by grants NIDDK 1R01DK129675 and NIDDK 1R01DK137473. We thank NYULH Microscopy Lab for consultation and the electron microscopy works. The Microscopy Lab is partially supported by NYU Cancer Center Support Grant NIH/NCI P30CA016087. We also would like to thank the Genome Technology Center at NYULH for RNA-sequencing. We thank the Histology core at NYULH for H&E and MTS staining.

## Author Contributions

Xiaozhong Xiong and Melody Ho contributed to this work equally in performing experiments, data analysis and preparation of manuscript. Karim Jaber, Rashmi Mishra, Amalya Charytan and Nadim Zaidan participated in animal work and paper preparation. Florencia Schlamp contributed to RNA-seq analysis. Lama Nazzal designed the study, supervised the experiments. Glenn Fishman and Lama Nazzal reviewed the manuscript.

**Correspondence** and requests for materials should be addressed to corresponding author Lama Nazzal.

## Data availability

All raw data of microarray analyses have been submitted to Gene Expression Omnibus (GEO) database (accession no. GSE297087). To review GEO accession GSE297087, go to: https://urldefense.com/v3/_https://www.ncbi.nlm.nih.gov/geo/query/acc.cgi?acc=GSE297087_;!!MXfaZl3l!cbtHZ9F0LzlAWM0MLQaeiP5l-mX1miasRccQFh6pxCGO2jcXl-TtJNjpEGXPiT8ux_PhR7RaMqU8wty8n2GLSQj_MshRUw$ and enter token **uloluemulhuhhev** into the box. Source data are provided with this paper. Other data that support the findings of this study are available within the article and Supplementary files.

