## Supplemental Figures for "Colonization with *Oxalobacter formigenes* slows the progression of CKD and reduces cardiac remodeling in CKD"

**Supplementary Material**

Supplementary Figures

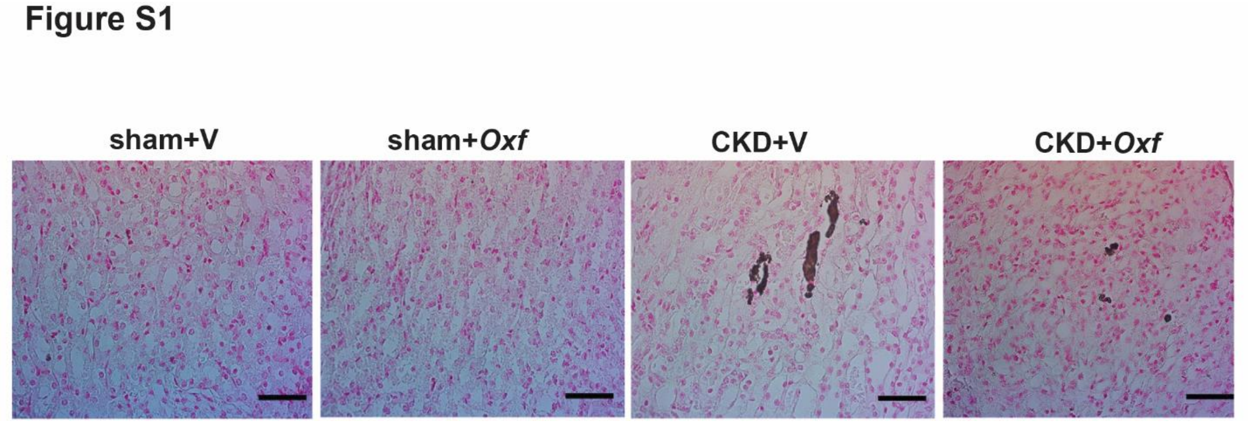


**Figure S1**. Pizzolato staining exhibited calcium oxalate crystals deposited in the kidneys of CKD mice.

### Figure S2

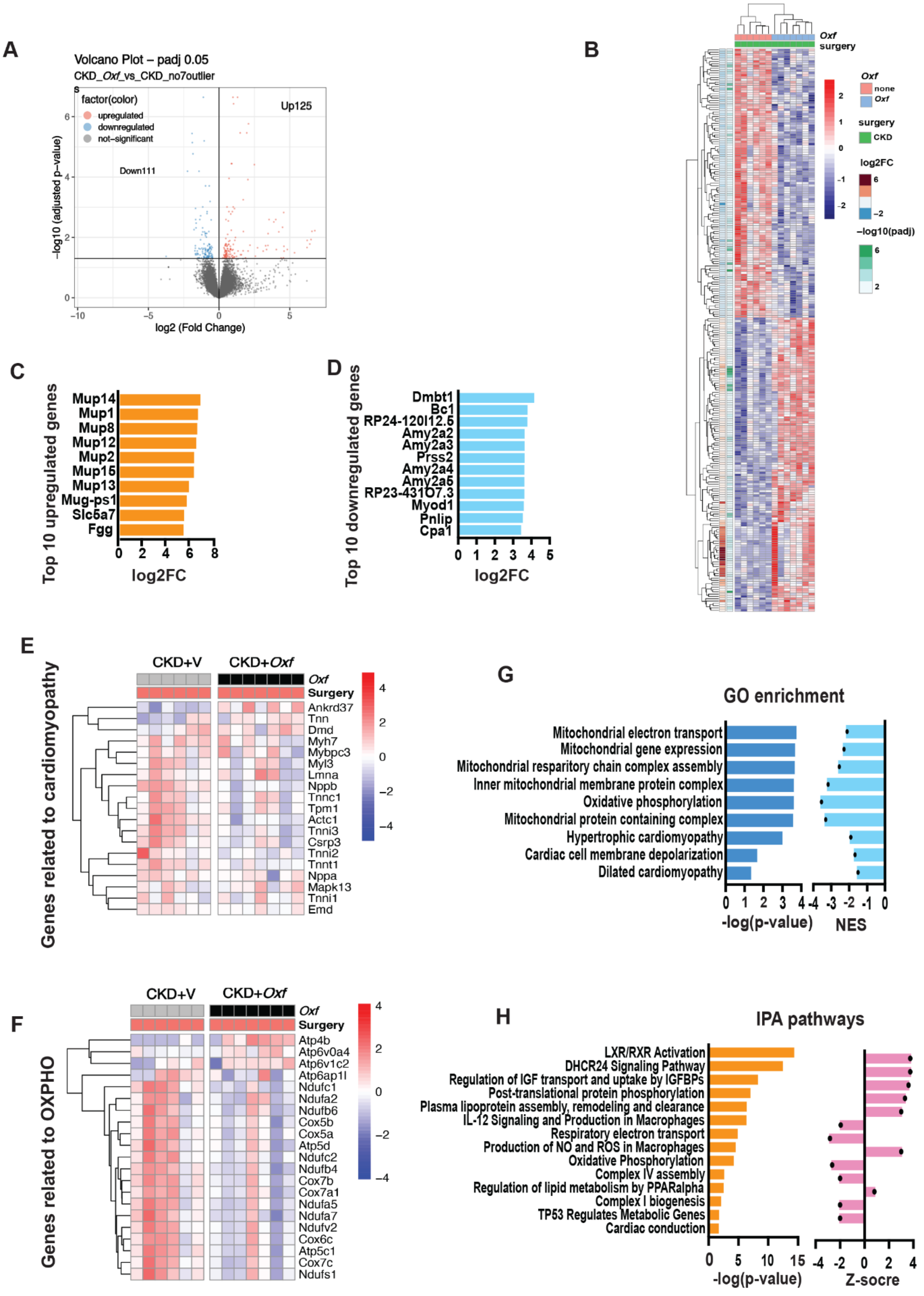
A
Volcano Plot – padj 0.05
CKD_Oxf_vs_CKD_no7outliers
factor(color)
upregulated
downregulated
not-significant
Up125
Down111
-log10 (adjusted p-value)
log2 (Fold Change)
B
Oxf
surgery
none
CKD
log2FC
-log10(padj)
C
Top 10 upregulated genes
Mup14
Mup1
Mup8
Mup12
Mup2
Mup15
Mup13
Mug-ps1
Slc5a7
Fgg
log2FC
D
Top 10 downregulated genes
Dmbt1
Bc1
RP24-120I12.6
Amy2a2
Amy2a3
Prss2
Amy2a4
Amy2a6
RP23-431O7.3
Myod1
Pnlip
Cpa1
log2FC
E
CKD+V
CKD+Oxf
Surgery
Genes related to cardiomyopathy
Ankrd37
Tnn
Dmd
Myh7
Mybpc3
Myl3
Lmna
Nppb
Tnnc1
Tpm1
Actc1
Tnni3
Csrp3
Tnni2
Tnnt1
Nppa
Mapk13
Tnni1
Emd
F
Genes related to OXPHO
Atp4b
Atp6v0a4
Atp6v1c2
Atp6ap1l
Ndufc1
Ndufa2
Ndufb6
Cox5b
Cox5a
Atp5d
Ndufc2
Ndufb4
Cox7b
Cox7a1
Ndufa5
Ndufa7
Ndufv2
Cox6c
Atp5c1
Cox7c
Ndufs1
G
GO enrichment
Mitochondrial electron transport
Mitochondrial gene expression
Mitochondrial respiratory chain complex assembly
Inner mitochondrial membrane protein complex
Oxidative phosphorylation
Mitochondrial protein containing complex
Hypertrophic cardiomyopathy
Cardiac cell membrane depolarization
Dilated cardiomyopathy
-log(p-value)
NES
H
IPA pathways
LXR/RXR Activation
DHCR24 Signaling Pathway
Regulation of IGF transport and uptake by IGFBPs
Post-translational protein phosphorylation
Plasma lipoprotein assembly, remodeling and clearance
IL-12 Signaling and Production in Macrophages
Respiratory electron transport
Production of NO and ROS in Macrophages
Oxidative Phosphorylation
Complex IV assembly
Regulation of lipid metabolism by PPARalpha
Complex I biogenesis
TP53 Regulates Metabolic Genes
Cardiac conduction
-log(p-value)
Z-socre

**Figure S2. RNA-seq analysis of genes and pathways in the hearts of CKD+*Oxf* vs CKD+V.** **A.** Volcano plot displayed genes differentially expressed (adjusted $p<0.05$). **B.** Heatmap of differentially expressed genes (adjusted $p<0.05$). **C.** Top 10 up-regulated genes (fold >2, $p<0.01$). **D.** Top 10 down-regulated genes (fold >2, $p<0.01$). **E.** Heatmap of differentially expressed genes related to oxidative phosphorylation (OXPHOS) ($p<0.05$). The color intensity represents log2FC changes. **F.** Heatmap of differentially expressed genes related to cardiomyopathy ($p<0.05$). The color intensity represents log2FC changes. **G.** GO enrichment of the significantly changed cellular components. Right side (blue) is -log(p-value), left side (light) is NES value. **H.** IPA pathways of differentially expressed genes (DEG) of the heart. Right side (orange) is -log(p-value), left side (pink) is Z-score score.
